# A non-invasive method for time-lapse imaging of microbial interactions and biofilm dynamics

**DOI:** 10.1101/2022.03.17.484708

**Authors:** Carlos Molina-Santiago, John R. Pearson, María Victoria Berlanga-Clavero, Alicia Isabel Pérez-Lorente, Antonio de Vicente, Diego Romero

## Abstract

Complex interactions between microbial populations can greatly affect the overall properties of a microbial community, sometimes leading to cooperation and mutually beneficial coexistence, or to competition and the death or displacement of particular organisms or sub-populations. Interactions between different biofilm populations are highly relevant in diverse scientific areas, from antimicrobial resistance to microbial ecology. The utilization of modern microscopic techniques has provided new and interesting insight into how bacteria interact at the cellular level to form and maintain microbial biofilms. However, our ability to follow complex intra- and inter-species interactions *in vivo* at the microscopic level has remained somewhat limited. Here, we detail BacLive, a novel non-invasive method for tracking bacterial growth and biofilm dynamics using high resolution fluorescence microscopy and an associated ImageJ processing macro (https://github.com/BacLive) for easier data handling and image analysis. Finally, we provide examples of how BacLive can be used in the analysis of complex bacterial communities.

**Importance:** Communication and interactions between single cells are continuously defining the structure and composition of microbial communities timely and spatially. Methods routinely used to the study of these communities at cellular level rely on sample manipulation what deprives from microscopic time lapse experiments on a given sample. BacLive is conceived as a method for the non-invasive study of the formation and development of bacterial communities, such as biofilms, and the dynamics of formation of specialized subpopulations in time-lapse experiments. In addition we prove that BacLive largely simplifies the analysis of the data generated.

## Introduction

Bacteria commonly live in structurally and dynamically complex biological systems forming surface-associated communities known as biofilms, which typically feature one or more bacterial species and/or other organisms in close proximity (1). Complex interactions at nutritional and chemical levels can confer a range of benefits including increased resistance to environmental stresses, metabolic exchange, better surface colonization, and increased horizontal gene transfer, which can greatly affect the final structure of the entire community (2, 3). Thus, cooperative or competitive interactions can lead to mutually beneficial coexistence, or to the displacement or death of particular organisms or sub-populations (4). In addition to single and multispecies biofilms, bacterial interactions between different biofilm populations are of interest for a wide variety of applications such as the study of antibiotic resistance mechanisms and development of new antimicrobial drugs (5–7). In these cases, bacterial populations are separated in the range of centimeters to millimeters, allowing long-distance effects to be studied.

Visualizing bacterial growth has been at the core of microbial science for over a century. Since the invention of the microscope in the 17th century, classical brightfield microscopy has long been used for the detection and classification of bacteria. Most of our understanding of bacterial interactions has been obtained through traditional microbiological techniques, which mainly address macroscopic changes visible to the naked eye (8). In recent years, light microscopy has been used to study whole bacterial populations, aiming to understand bacterial interactions and/or biofilm formation dynamics at cellular and subcellular levels (9). The development of bacterial communities was initially documented using standard digital cameras that permit the capture of macroscopic 2D images of colonies grown on solid culture media (8, 10, 11). Changes in colony attributes were followed over time by placing inoculated plates in incubators for several days and imaging at different timepoints during the experiment (12, 13). This simple methodology is ideal for experiments where only limited changes occur over time, limiting dehydration and the potential problems caused by condensation, as plates remain in optimal growth conditions for most of the experiment. However, these methods are limited in the study of more complex biological processes or microbial interactions. More sophisticated imaging approaches have been developed to deal with these issues. The MOCHA (microbial chamber) method (10), also based on macroscopic imaging, consists of a sealed chamber with built in humidity control, and crucially, houses the camera inside the chamber, and is all controlled by an external computer (Table 1). With this method, Petri dishes are prepared with paper wicks connected to a water reservoir to maintain hydration of the culture medium. Altogether, these improvements allow for the imaging of bacterial growth for long periods (up to 40 days) while monitoring environmental conditions inside the chamber. A similar approach was developed by V. M. Zacharia et al. (11) to perform time-lapse imaging using a fluorescence microscope and a USB-controlled heated mug warmer with a temperature sensor to monitor real-time temperature within the chamber and prevent condensation.

**Table 1.** Comparison between 2D and 3D approaches used to visualize bacterial growth and their interactions.

| Method | Monitoring time | Resolution | Invasiveness | Continuity | Reference |
| --- | --- | --- | --- | --- | --- |
| 2D imaging with standard camera in incubators | Up to 4-7 days | Centimeter to millimeter scale | No | No | Traxler et al., 2013 |
| 2D imaging with standard camera in MOCHA | Up to 40 days (without contamination) | Centimeter to millimeter scale | No | Yes | Penil-Cobo et al., 2018 |
| 3D imaging by cryostat sectioning | Up to 4-7 days | Micrometer scale | Yes | No | Vlamakis et al., 2008 |
| 2D Time-lapse confocal microscopy | Up to 1 week | Micrometer scale | No | Yes | Nadezhdin et al., 2020 |
| 3D Time-lapse confocal microscopy | Up to 1 week | Micrometer scale | No | Yes | Molina-Santiago et al., 2019 |

All of the above imaging methods are macroscopic in nature and lack the resolution to study bacterial interactions at the cellular level. The classical method based on mounting bacterial cultures between slides and coverslips is excellent for counting bacteria, visualizing cellular morphology and other bacterial properties (14–16) but it is not useful for colony progression studies, given the loss of all spatial and structural information. In recent decades, the development of new microscopes and the increasing importance of microbial ecology and biotechnology have led to the establishment of a variety of strategies for the study of microbial interactions and community development at the microscopic scale. For example, cryostat sectioning has been used with fluorescently-labelled bacteria to demonstrate the existence of heterogeneous populations dedicated to different physiological functions and follow changes during *B. subtilis* biofilm development (17) (Table 1). However, it is an invasive and destructive method. Its temporal resolution is limited to the number of colonies frozen and sectioned, making it difficult to link sequential events that may vary between individual colonies. Another approach consists in combining high resolution fluorescence microscopy with bacteria expressing different fluorescent reporters to characterize spatial expression patterns in colonies of *Bacillus subtilis* (18). This approach consists of cutting a piece of agar containing the colony edge and flipping it onto a glass-bottomed dish before imaging, thus allowing visualization of reporter construct expression patterns at different levels of the bacterial colony (Table 1).

In order to study interactions between bacterial biofilm populations on solid media, we have developed a method that combines microscopic resolution, 3D acquisition, differential labelling and continuous imaging for up to 5 days (14). This approach takes advantage of high quality long working distance immersion optics combined with glass-bottomed Petri dishes to visualize cellular dynamics within growing colonies, in a truly non-invasive manner (Table 1; Figure 1A). Here, we describe the acquisition methodology step-by-step, explaining the rationale behind it and its main advantages and disadvantages. We also detail our post-acquisition methodology, which includes a freely available data handling tool (https://github.com/BacLive/) that aids efficient processing and analysis of the results generated by the BacLive method. Finally, we show specific examples of how this methodology can be used in: i) the study of bacterial biofilm interactions on solid media, and ii) dynamics of the formation of specialized subpopulations within a *B. subtilis* biofilm.

**Figure 1.**
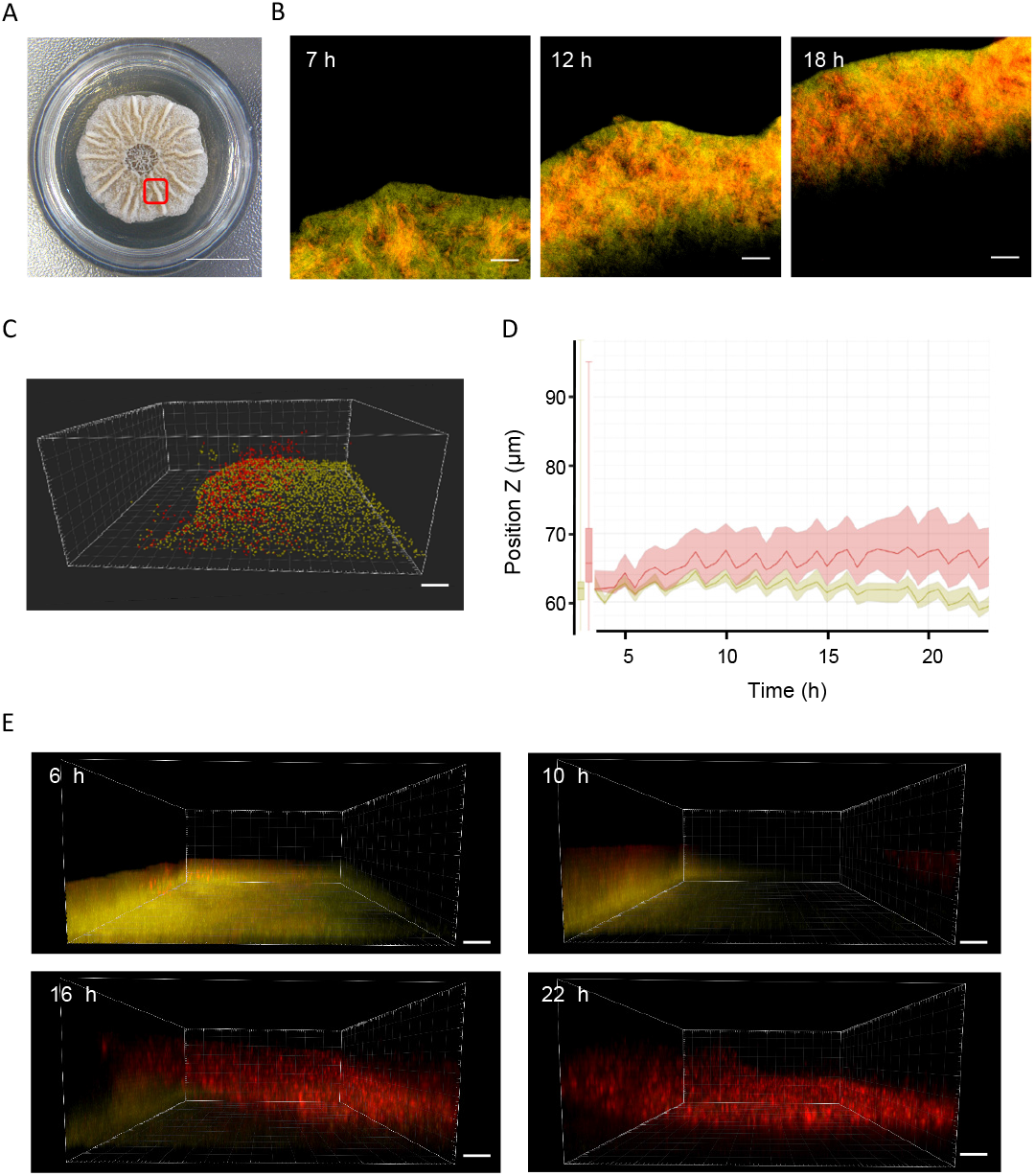
Schematic representations of BacLive and plate position in an inverted fluorescent microscope. A) Schematic representation of the main steps to perform bacterial interactions and biofilm development assays using confocal microscopy and Bac-Live processing image. Step 1 highlights the most important points for sample preparation. Step 2 indicates main concerns related with microscopy setup and conditions, including multiposition acquisition, growth conditions and technical setting as number of slices in a Z-stack and image dimension. Step 3 focus on data analysis using Bac-Live as the main tool for data image processing using FiJi and more complex analyses using commercial softwares such as Imaris. B) Schematic representation of the plate position in an inverted fluorescent microscope in which are highlighted the main elements that need to be taken into account for a correct setup, such as the need of immersion media, objective lens characteristics, position of colonies for bacterial interaction and the use of 35 mm glass-bottomed Petri dishes.

## Results and Discussion

### Acquisition Methodology

Confocal laser-scanning microscopy (CLSM) has emerged as one of the dominant methodologies for fluorescence bioimaging over the last 30 years (reviewed by Pawley, 2006). In the context of biofilm imaging, a confocal system provides two main advantages over widefield ones: firstly, they are more robust at optically reducing extraneous fluorescence, agar auto-fluorescence and reflections, and secondly, they provide higher resolution 3D information of colony and biofilm development. On the other hand, CLSM systems have some disadvantages for bacterial imaging. Firstly, they tend to be relatively insensitive compared to widefield systems. Moreover, they are typically associated with higher photobleaching and phototoxicity. However, on balance, we have found that background fluorescence, reflections and bacterial autofluorescence were more significant issues than pure sensitivity for biofilm imaging. Even performing long time-lapse studies using 10x and 25x objectives, we did not observe significant photobleaching or phototoxicity, although at higher magnification levels this is more likely to become an issue. Another limitation of point scanning image acquisition is that it can be slower than CCD/CMOS-camera based systems. Long-term biofilm experiments do not require particularly high-speed imaging, and would otherwise generate prohibitive amounts of data. In addition, most CLSM systems can capture multiple fluorescence channels simultaneously, whereas widefield systems usually need to capture each channel separately, meaning that the real-world difference in speed is often minimal. It is important to point out that while here we focus on a confocal-based approach, the basic methodology is compatible with a variety of other modalities including brightfield, traditional widefield fluorescence and deconvolution-based approaches, although the post-processing and analysis would specifically differ in each case.

### Microscope type (inverted vs upright)

Research microscopes are arranged in either upright or inverted configurations, where the objective lens is located above or below the sample, respectively. Figure 1B shows how bacterial growth could be followed in the inverted configuration, which has been successfully used in our experiments, but upright systems should work well and may have some advantages. Using a whole microscope heated enclosure, we did not have significant problems with condensation build-up. However, this potential problem would be reduced using an upright system, where the culture media is inverted and excess condensation would not drip onto the agar surface.

### Glass-bottomed dishes

Most research microscope objective lenses are optimized for coverslip thicknesses of 170 microns with a tolerance of +/- 20 microns (type #1.5 coverglass). Focusing through different thicknesses of glass or materials with different properties can reduce resolution, sensitivity and other optical artifacts. Therefore, for optimal results we used 170 +/- 10 micron coverglass-bottomed Petri dishes (Figure 1B). It is worth noting that some objectives have correction collars that can compensate for different coverslip thicknesses and also that low magnification non-immersion objectives will normally be less sensitive to suboptimal optical conditions, and may allow visualization through standard plastic-bottomed Petri dishes. However, given the already challenging optical path traversing the agar, we would strongly recommend avoiding this option if possible.

### Temperature control

For most bacteria, temperature control is a crucial factor in colony growth, biofilm formation and their interactions with other organisms. Therefore, it is crucial to control the environmental conditions as much as possible (Figure 1A). For long term imaging, maintaining temperature is also critical for optical stability. Where possible we recommend the use of a full microscope enclosure with temperature control and a temperature sensor positioned as close as possible to the Petri dish. Thus, variations that could affect bacterial growth or optical stability are reduced. Enclosure-based heating should also reduce condensation problems compared to a plate/stage heating solution.

Conditions will vary greatly depending on the species of bacteria being studied and the experimental goals. In most cases, experimental temperatures will range from 28 °C to 37 °C. In practice, temperature control is easier the greater the difference from the ambient temperature. Both the microscope excitation laser (or fluorescence lamp) and microscope electronics will contribute to a locally increased temperature close to the sample. In experiments with an ambient temperature of 21 °C, we routinely measured temperatures adjacent to the sample of 26-27 °C during long time-lapse experiments without any additional heating. Therefore, it would be difficult to perform experiments at lower temperatures without lowering the ambient temperature or some kind of active cooling. This emphasizes the importance of regulating temperature based on measurements that are as close to the sample as possible.

### Agar composition and thickness

One of the key innovations in our methodology is the idea of imaging colonies through the agar substrate. This means that immersion objectives can be used to visualize biofilm dynamics with a greatly improved resolution versus non-immersion objectives. It also means that imaging can be performed on a closed Petri dish without a loss of imaging quality. The crucial limiting factor is the working distance of the objective lens. Most high-quality oil objective lenses are only capable of focusing up to 90-200 microns into a sample, which would make it impossible to focus through a reasonable volume of solid substrate. An increasing number of high-quality, long working distance (LWD) objectives are being offered by microscope manufacturers. Most commonly, these lenses are designed to use water as an immersion medium. This improves imaging quality when imaging through water-based substances, which would typically be cell-culture medium or animal tissues, but should also hold true when imaging through a water-based solid medium. For our work, we took advantage of a Leica 25x water immersion objective (HC FLUOTAR L 25x/0.95 W VISIR) with a free working distance of 2.4 millimeters. The objective has a relatively high numerical aperture (NA) meaning a theoretical maximum xy resolution of 205 nanometers, which should be more than sufficient for individual bacterial imaging in a confocal setup, depending on scan conditions. The free working distance of the objective determines how much solid media can be used while still allowing us to focus on the nascent colony at the start of the experiment. We tested the ability of different bacterial strains to grow on different quantities of Lysogeny-Broth (LB) and Msgg solid medium and found normal growth occurred with as little as 1.3 mL of solid medium in a 35 mm diameter Petri dish with a starting thickness of 1 mm. However, normal growth was affected if volumes were reduced below 1.3 ml (Suppl. Figure 1). These values represent a good starting point but we would always recommend early testing of this aspect as media volume sensitivity may vary depending on the bacterial species, mutations and composition of media used. If the minimal media volume results in a greater thickness than the free working distance of your objective(s), then an alternative imaging strategy would be required.

### Objective lens choice and troubleshooting

Evaporation of the water immersion media is a major problem for long-term time-lapse imaging. Typically, after about 2 hours, the experiment would have to be discontinued to add new water immersion media, which is impractical in multi-day time-lapse imaging. For many microscope/objective combinations, automatic water dispensers are available that replenishes water at the objective lens throughout the experiment. As this kind of dispenser was not available in our case, we took an alternative approach, using a special silicone oil-based immersion medium (Immersol W 2010 Immersion Oil, Carl Zeiss), with a refractive index of 1.33 suitable for water immersion objective lenses. We found that this method worked very well with our Leica objective lens for this kind of long-term experiment. As oil does not have the same surface tension as water, there is a limit to how much oil can be added without it dripping down from the lens-dish interface. Fortunately, as the objective is relatively close to the base of the Petri dish (i.e. at or near to the limit of its focal distance), only a relatively small amount of immersion medium is required. For multi-position experiments (see below), best results were obtained when oil was spread across the glass bottom, in addition to oil being added directly to the objective. By doing so, we were routinely able to image bacterial dynamics, uninterruptedly, over multiple days.

### Methods for dealing with evaporation and media shrinkage

Our methodology allows for high quality imaging while maintaining the Petri dish closed, which helps minimize evaporation. However, especially in a heated environment, evaporation is inevitable. Evaporation changes the relative position of the colony with respect to the objective lens, so the colony would gradually drift out of focus after a short period (Figure 2A). Evaporation could be reduced using methods similar to those employed by the MOCHA method (M. Penil Cobo et al. (10). Alternatively, an autofocus method based on fluorescence intensity or contrast could be employed to follow colony position changes. For our methodology, we chose to use an approach based on the acquisition of large 3D volume(s) that cover the future positions of the colony. We discovered that colony movement due to evaporation and media consumption was highly consistent and predictable, with a constant movement over time of approximately 2.5 microns per hour (Figure 2B) with negligible differences observed between different regions of the same plate (Figure 2B zoom). Thus, it is possible to calculate the total volume needed to follow colony dynamics over the planned length of the experiment (Suppl. Figure 2A). This “brute force” approach meant that we were not dependent on specific fluorescence expression to ensure correct autofocusing. Furthermore, it also ensures that we could follow biofilm dynamics in 3D without having to know in advance how colony development would proceed. The main limitation to this approach comes from data management and system stability. Data volumes can be reduced by taking advantage of the predictability of colony movement and performing acquisition in sub-volumes to cover different temporal periods as illustrated in Suppl. Figure 2B. Many microscope systems allow this kind of more complex acquisition pattern to be programmed automatically, but could also be implemented manually every 24 hours, for example. In practice, even with a 10-year old Leica SP5 CLSM system, we could routinely acquire several plate positions over multiple days generating very large Z-series with final data volumes exceeding 20-30 gigabytes. However, this approach does bring a number of considerations and data processing issues as we discuss below.

**Figure 2.**
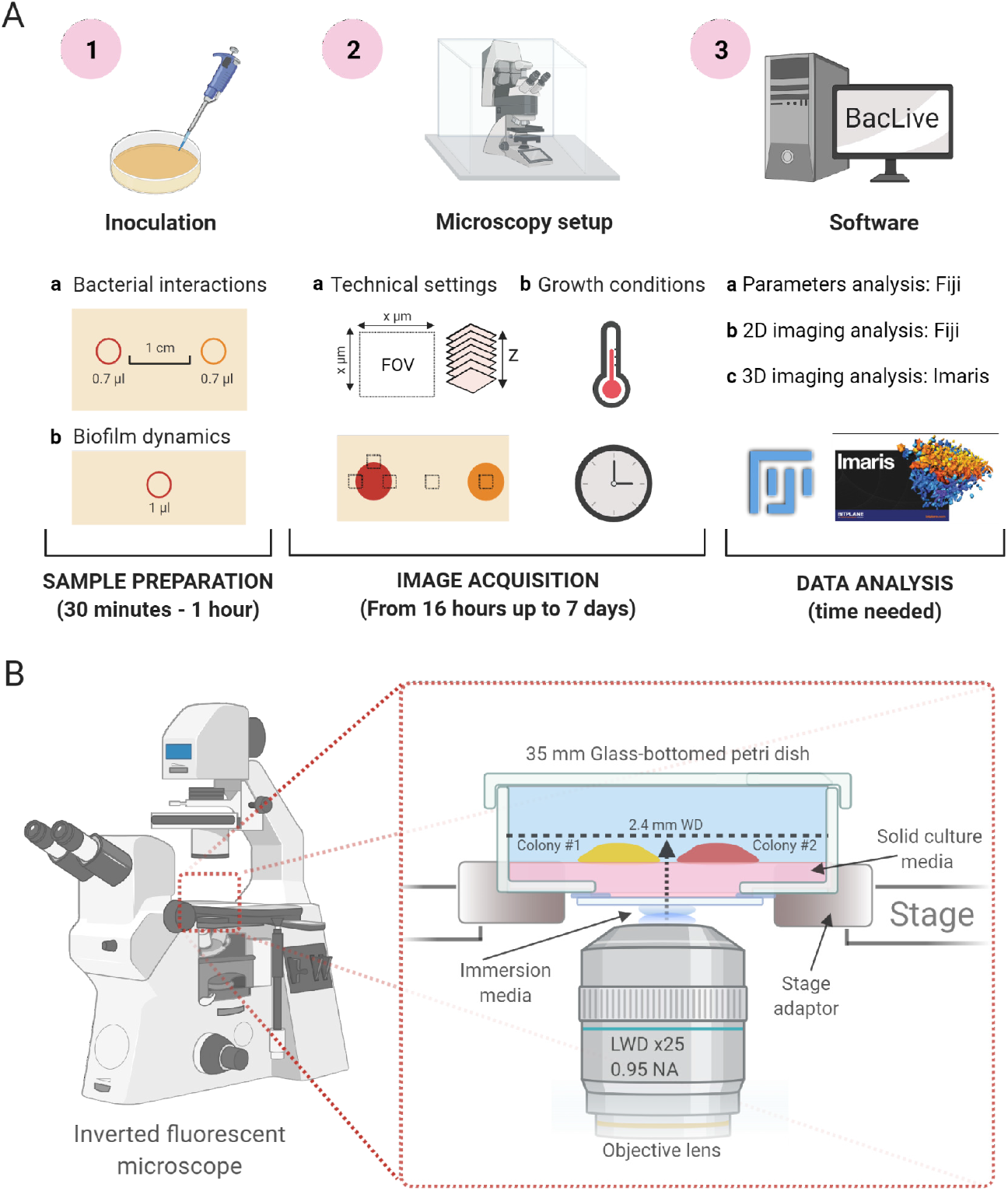
Technical settings for experimental processing and data analysis. A) Evolution of solid media thickness along the experiment. Due to evaporation, the solid media experiences a thickness reduction. B) Colony Z-position changes occurring in a timelapse (72 h) pairwise interactions experiment (strains labelled with CFP and RFP). Zoom for the first 24 h show slight differences between fluorescent strains but, overall, the evaporation is continuous during the whole experiment making the movement calculation predictable. C) Schematic representation of a multiposition experiment. Each square represents one image-capture position. We usually select edge positions, inner colony positions and expected contact positions. D) Comparison of unprocessed and post BacLive processed image data shows the differences according to the number of slices and data size reduction. The number of slices is dramatically reduced due to the filtering and selection of slices with fluorescence.

### Acquisition Frequency / Imaging conditions

For a 3D time-lapse experiment, the distance between acquisition slices, or Z-sampling frequency, determines the 3D resolution and has a critical effect on data size and acquisition time. Thus, it is often necessary to sacrifice 3D resolution to allow for a reasonable acquisition speed. We would typically set the Z-sampling interval between 1 and 3 microns depending on the experiment. Given that the axial resolution of our objective is ~550 nm, our datasets are considerably undersampled, optically speaking, but are still sufficient to provide valuable 3D information about colony dynamics while limiting acquisition time and dataset size.

Imaging conditions are another major consideration where image quality, sensitivity, laser power and imaging speed must be balanced. It will normally be necessary to perform pilot experiments to establish appropriate conditions as the expression of fluorescent proteins can vary greatly between different constructs, strains and at different stages of colony growth. In the example described in more detail below, each multichannel 2D image or Z-slice had an acquisition time of ~3 seconds. For a 40-hour time-lapse experiment we predicted a Z drift of ~100 µm but choose to acquire a larger 250 µm volume in case we wished to extend the experiment beyond the initial 40 hours. We combined this with a ~2 µm Z-stack sampling frequency, equating to a total of 128 slices and ~6-minute acquisition time per timepoint. We established a 30 minute interval as being sufficient to follow bacterial growth and biofilm dynamics, thus capturing 84 timepoints and a total of 32,256 images (2 fluorescence channels and transmitted light).

### Multi-position acquisition

For our studies of biofilm dynamics in the interaction between *Pectobacterium carotovorum* and *Bacillus amyloliquefaciens* FZB42 (See ‘ study of bacterial population dynamics in interactions on solid media’ section below) we wanted to be able to follow changes of each colony and their interaction during the same experiment. In order to do this, we took advantage of the motorized stage and multi-point acquisition, a feature common to many mid-to-high end research microscopes. At the start of the experiment, we selected stage positions corresponding to different colony regions and regions where we predicted colonies might come into contact. We then defined Z-stack positions for each region separately, as the agar surface is usually not exactly level with respect to the microscope, but otherwise defined them using the same total volume and Z-sampling frequency (Figure 2C). Multi-position time-lapse acquisition proceeds by acquiring each Z-stack in turn. The time-lapse interval defines the period between starting the first Z-stack of each timepoint, thus all acquisitions must be completed within the time-lapse interval. This ensures that all fields have the same imaging interval even though they are not simultaneous. Acquiring multiple microscope fields also multiplies all of the limiting factors in terms of balancing image quality and imaging time. Reducing image resolution from 1024 × 1024 to 512 × 512 was typically necessary to avoid excessively long time-points and, especially, to avoid generating prohibitive amounts of data.

As we have seen, a large number of variables need to be balanced in terms of bacterial inoculation, culture media and acquisition settings to optimize a time-lapse experiment. Different bacterial models and experimental priorities will require different compromises to be made for practical reasons. However, as with any time-lapse experiment, it is essential to validate the imaging configuration with appropriate controls to monitor the extent that imaging conditions might alter bacterial and biofilm development. If incubator grown control plates show different patterns of growth or interaction then it may be necessary to further optimize environmental conditions or alter imaging conditions (e.g., reduced laser power). In some cases, the development of improved reporter constructs with enhanced expression levels or better stability, may help achieve better results.

### Image Processing Methodology

#### Data management (in general) / Data Compression in situ

The primary disadvantage and limiting factor of the strategy we have implemented in this method is the very large amounts of data that are typically generated. Figure 2D shows an example where a 73-hour acquisition generated 27.4 Gb of data per position. We routinely used this method to acquire data from 6 different positions over 5 days. Given a typical plate z-drift of ~2.5 μm per hour, this requires a minimum of ~300 μm total Z-stack thickness and 150 image sections given a 2 μm z-spacing. In the example shown below, we acquired two fluorescence channels for CFP and GFP plus brightfield. Using a 512 × 512 XY resolution and 8-bit images this is equivalent to 0.66 Gb per time-point or ~158 Gb over 5 days with a 30-minute acquisition interval. Therefore, finding efficient ways to deal with these large datasets is the main focus of the remainder of the article.

Depending on the local infrastructure and faculty organization, it may be necessary to copy data from the acquisition computer before processing. To make this procedure more efficient, we recommend saving each acquisition field of view as a separate file. It is much easier to deal with several 5 Gb files than a single 30 gigabyte one. Secondly, if the microscope platform saves in a non-compressed format, files can be compressed by as much as 80% using open-source software such as 7zip (https://www.7-zip.org/). This can make the job of copying and storing large data files much easier, although it may increase the risk of data corruption during the copying process. We would recommend not deleting the original uncompressed data until it is safely copied and verified elsewhere. If your local environment permits processing data *in situ* as described below, it may be a preferable option.

#### BacLive data preprocessing ImageJ/FIJI macro

By acquiring large Z-stacks that cover the predicted trajectory of colony growth, media consumption and media shrinkage, it is inevitable that the vast majority of the image slices collected will contain non-useful data. Our objective is to trim this excess data while compensating for colony movement over time, thus allowing us to follow biofilm dynamics and colony interactions. Our strategy to do so is based on the observation that under our acquisition conditions, colony movement is linear over time (Figure 2B). Therefore, by knowing the agar surface starting and ending positions, we can calculate the average change in Z position over time in terms of Z-slices. Using this value, we aim to process the full dataset using a “floating window” crop of Z-slices that follows the agar surface / colony position (Figure 2D).

This relatively simple strategy is surprisingly effective in terms of compensating for colony movement/media shrinkage. However, it is still very time consuming when performed manually for each time-point. To make this process more efficient we have devised a macro script written for the ImageJ/FIJI open-source image processing packages (20, 21). The script (freely downloadable from https://github.com/BacLive/) assists the user in detecting the average Z-slice change value and then applying a floating window crop and generating a processed 3D time-lapse dataset, where the plate surface should remain relatively fixed in the Z-plane and eliminating most of the unneeded Z-slices. As ImageJ/FIJI supports a wide range of image formats through the Bioformats plugin (22), the same methodology should be effective regardless of the microscope type used for acquisition.

Over time our BacLive processing macro has developed considerably, offering a relatively simple and straightforward interface for image processing, that is designed to avoid some of the memory limitations that can be problematic when dealing with large time-lapse dataset. For example, it is now compatible with virtual datasets opened with the Bioformats plug-in, making it relatively painless to process huge datasets without needing specialist workstations with large amounts of RAM. Processing is essentially divided into two sections as shown in the BacLive workflow (see Suppl. Figure 3). Firstly, it gives the user the option of generating a graph showing average image intensity by channel, Z-position and time. This will aid in choosing appropriate timepoints and channels for subsequent steps for tracking plate surface movement. Next, BacLive calculates the rate of surface movement (or Displacement Value) based on automatically-calculated maximum intensities or based on user-selected best focus slices. Start and End positions can be any period where the plate surface/biofilm can be visualized, thus it is not necessary for the biofilm to be present in a given field-of-view for the whole experiment. The manual slide selection method is also compatible with brightfield images and it is usually the best option when fluorescence is very weak. The automatic method is based on fluorescence intensity and is designed to set the central Z-position at the point with the highest intensity. Again, the user can choose specific reference timepoints for this calculation and the macro will extrapolate to allow processing of the whole image stack.

The second processing phase (Suppl. Figure 3) is performed using the calculated Displacement Value to apply a floating Z-window crop progressively over time and follow the plate surface position with a user-specified thickness in slices. The software will try to estimate the starting crop position based on manual or automatic values but users can also manually specify both starting position and Displacement Value. During processing, the macro writes timepoints as temporary files (.tif) to avoid memory issues, before reassembling the reduced dataset with its original LUT (color) values and calibration. The benefits of BacLive can clearly be seen in Figure 2D, where a 27.4 Gb dataset can be quickly reduced to a much more manageable 3.4 Gb one, without losing the 3D interaction dynamics of the two colonies.

#### Study of bacterial population dynamics in interactions on solid media

To illustrate how the BacLive imaging and processing methodology (resumed in Figure 1A) can be used for the study of practical problems in biofilm dynamics and bacterial interactions, we have focused on two examples: i) bacterial interactions between *Pectobacterium* and *Bacillus* species, and ii) the expression pattern and spatial distribution of different cell types in a single *Bacillus subtilis* biofilm.

The first example focuses on interactions between two ubiquitous soil species populations: *Pectobacterium carotovorum*, a phytopathogenic species that produces and secretes cell wall degrading enzymes leading to rotting and decay of their plant hosts (23), and *Bacillus amyloliquefaciens* FZB42 (referred as FZB42), a plant beneficial bacteria with antibacterial and antifungal activity against a wide range of plant pathogens (24). Initially, pairwise interactions on LB plates for 48 h between *P. carotovorum* and FZB42 showed the inhibition of *P. carotovorum* growth in the interaction area, where an inhibition halo was formed (Figure 3A). To study this effect in more detail, we analyzed interaction dynamics using the BacLive method. The FZB42 strain was fluorescently labelled with CFP, while *P. carotovorum* was labelled with GFP, as described in the Materials and Methods, and spotted on LB solid media at a distance of 0.7 mm on glass-bottomed Petri dishes. Temperature was controlled and maintained at a constant 28 °C during the experiment. Several Z-stacks at 6 different positions were recorded over 24-48 h to analyze the leading edge of each colony and the predicted zones of physical contact between the colonies (Suppl. Figure 4A). After data acquisition and image processing using the BacLive macro, we could analyze and quantify colony interactions at the different positions monitored. Initially, both bacterial colonies were growing separately and advanced at constant rates (Figure 3B-D). Time-lapse images at two positions (Figure 3C and 3D) allowed us to calculate the expansion rate of *P. carotovorum* at 40 µm/h and FZB42 at 100 µm/h after 8 h of growth (Suppl. Figure 4B). Both colonies reached one of the intermediate acquisition zones after 15 h of growth. Here, we observed the arrest of *P. carotovorum* colony growth while FZB42 continued its movement towards *P. carotovorum* (Figure 3E and Suppl. Movie 1). Measurement of colony movement confirmed this observation: FZB42 advanced at a speed of 10 µm/h, while *P. carotovorum* had initially stopped its advance, and then retreated at a rate of 5 µm/h (Figure 3B). The data collected allowed us to measure the distance between the two leading edges, showing that, at the moment the two colonies were recorded in the intermediate area, they were 160 µm apart and this distance progressively diminished until the colonies were practically in contact (Figure 3B). These results are surprising since the macroscopic view of the interaction (Figure 3A) showed a clear inhibition of *P. carotovorum*. However, our microscopic time-lapse data indicates that a viable *P. carotovorum* population is still present in the leading edge of the colony and this results in cell-to-cell contact with FZB42. In order to get a deeper understanding of the interaction, we took advantage of the 3D Baclive dataset and analyzed evolution of the two populations using Imaris software. From 18 h and 36 h of interaction, we could clearly observe the death of *P. carotovorum* at the time of continuous FZB42 advancement. Interestingly, when the two colonies came into contact, their leading edges became thicker (Figure 3F and Suppl. Movies 2 and 3). This suggests specific short-range interactions between the two populations. By combining BacLive with mutational studies, in the future, it may be possible to understand the molecular basis of these interactions. Our results show how this non-invasive technique combining fluorescence labelling with long-term three-dimensional acquisition can provide unique insights into the complex interactions between different bacterial populations.

**Figure 3.**
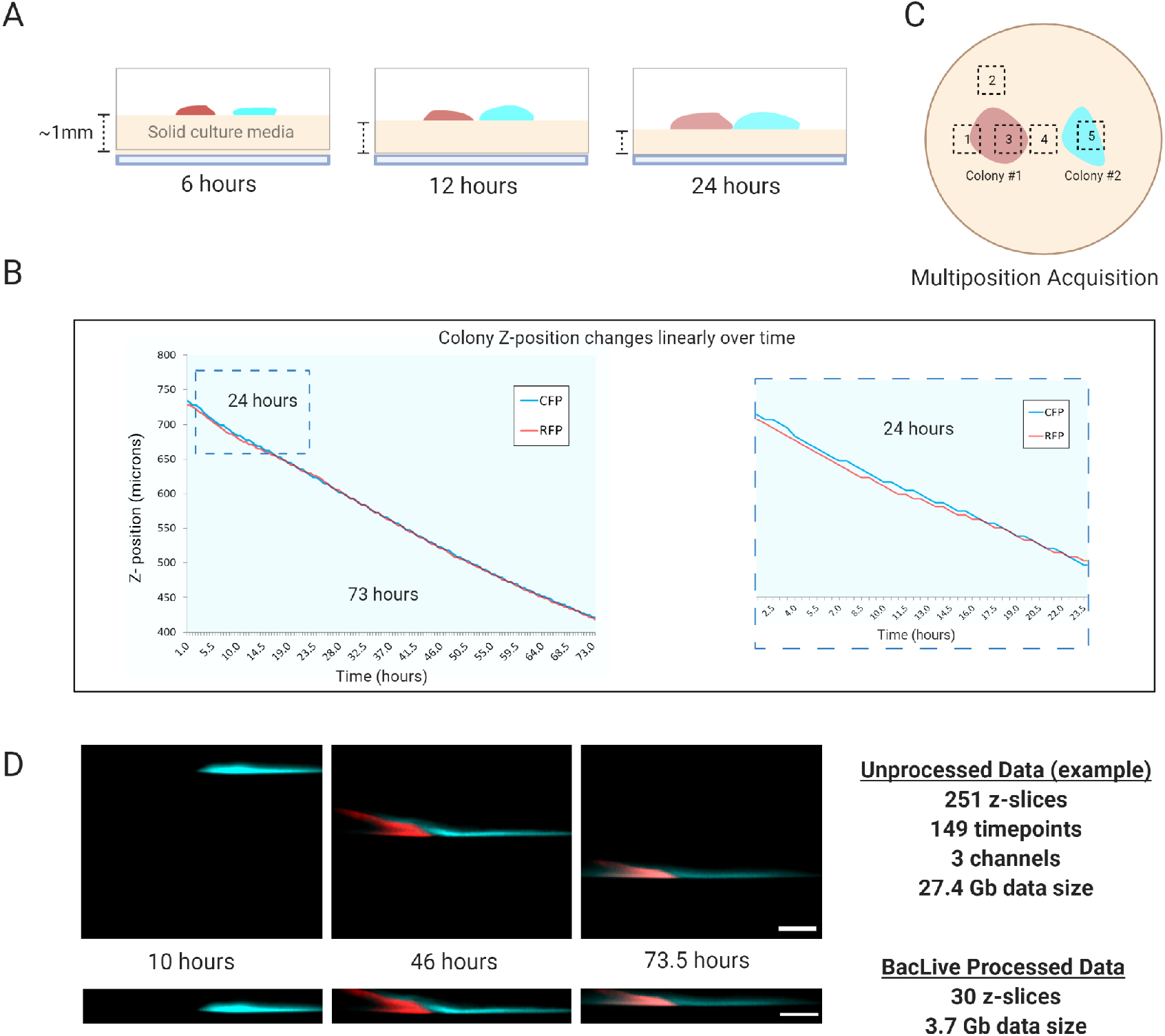
FZB42 inhibits the initial growth of *P. carotovorum* and they grow in a wall-like structure when get in contact. A) Pairwise interaction between FZB42 and *P. carotovorum* shows inhibition of *P. carotovorum* colony from a macroscopic perspective. Scale = 1 cm. B) Expansion rates of the FZB42 and *P. carotovorum* leading edges and distance between both strains in the long-term stage of the interaction (18-36 h). The blue line represents the FZB42 leading edge, and the green line represents *P. carotovorum* leading edge. The yellow area represents the distance between the two populations during this stage of the interaction. C) 2D images and Z-project representations of the *P. carotovorum* growth in the short-term timeframe of interaction (2 and 4-5 h). D) 2D images and Z-project representations of the FZB42 growth in the short-term timeframe of interaction (5.5 and 9 h). E) 2D images and Z-project representations of the interaction area where *P. carotovorum* and FZB42 get in contact (18 and 36 h). F) 3D surface representations of the interaction area show the initial contact between *P. carotovorum* and FZB42 and the FZB42 advancement. In addition, both colonies grow in a wall-like mode when they get in contact. Scale = 50 um.

#### Analysis of bacterial subpopulations during the *Bacillus subtilis* biofilm formation

Several studies have described the division of labor occurring in a bacterial colony during biofilm formation (18, 25). However, as mentioned above, the use of imaging technology for the study of subpopulations has typically required invasive sample processing, which typically risks deformation of colony morphology, and limits the study to specific time-points (17). With the use of the BacLive methodology described in this manuscript, we can follow the evolution of biofilm formation and the initiation of bacterial subpopulations and their development at the cellular level. To exemplify this, we have used a *B. subtilis* biofilm as a model to evaluate the timing and spatial distribution of two subpopulations: i) cells expressing TasA, a protein involved in the correct formation of the extracellular matrix (15, 26, 27); and ii) colony expansion based on *motA* expression, required for flagellar motor rotation. Promoters of both genes were placed upstream of sequences encoding two different fluorescent proteins (P_*tasA*_-mcherry and P_*motA*_-YFP), and integrated in two different neutral loci, *lacA* and *amyE*, respectively. 0.7 µl of the resuspended bacterial solution was used to inoculate Msgg solid media and the experiment was performed as described in the previous section (also see the Material and Methods section). We recorded a total of 6 different plate positions in order to analyze changes at the colony border and more central parts of the colony (Figure 4A). In addition, two extra recording positions were placed to capture later stages of *B. subtilis* colony expansion. Images were initially processed with BacLive, and then with Fiji and Imaris for further processing, visualization and analysis. When analyzed as a 2D projection, we could obtain a preliminary idea about the timing of promotor expression, however following the exact localization of motile cells and TasA producers was not easy (Figure 4B and Suppl. Movie 4). As expected, promoter expression in the inner colony gradually disappeared due to technical and microbiological factors. We next analyzed biofilm formation in 3D using IMARIS, which allowed us to evaluate the relative position of the two bacterial subpopulations with respect to the agar surface. For that, using IMARIS, we identified fluorescent bacteria foci as spots and measured their distance from the agar surface over time using the 3D BacLive-processed data at each timepoint (Figure 4C and Suppl. Movie 5). Figure 4D shows the average Z-position (height) of the two sub-populations with respect to the agar surface over time. The results clearly show that cells expressing TasA are mostly located in the upper region of the colony involved in extracellular matrix formation while cells expressing MotA are in closer contact with the solid media, thus permitting colony expansion. Significantly, one of the positions selected for time-lapse analysis coincided with a site of wrinkle formation. Wrinkles are structures typical of those formed in a *Bacillus subtilis* biofilm and are known to be empty structures permitting the movement of water and nutrients inside the colony (28). Interestingly, our time-lapse analysis showed the role of motile cells and TasA producing cells in wrinkle formation (Figure 4E and Suppl. Movies 5 and 6). While motile cells were associated with initial colonization and the expansion of colony borders, they did not appear to coincide with wrinkle formation. On the other hand, we observed TasA expressing cells adopting a distribution coinciding in time and space with wrinkle formation (Figure 4E).

**Figure 4.**
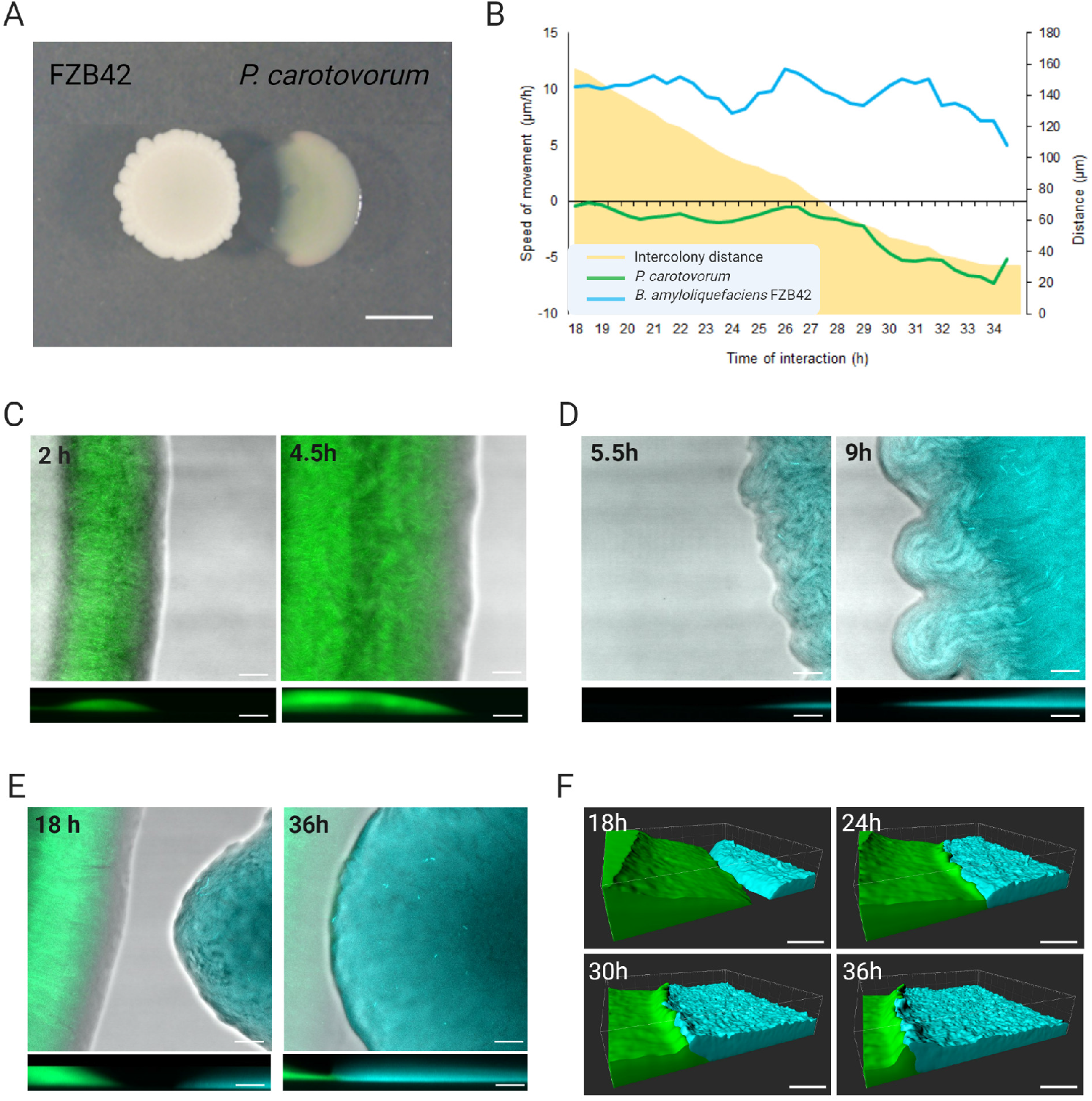
TasA expressing sub-population grows in the upper region of the biofilm while motility cells expand *Bacillus* colony in the contact with the culture media. A) *Bacillus* colony grown in Msgg culture media exhibiting its typical biofilm growth. Red square indicates one of the regions analyzed with the presence of wrinkles. Scale = 1 cm. B) 2D images representing both sub-populations during the colony growth (TasA expressing cells are fluorescently labelled with mcherry (red) while motility cells expressing MotA are labelled with YFP (yellow)). Scale = 20 µm. C) 3D image of a *Bacillus* colony growing on Msgg. Each spot represents cells expressing TasA (red) and MotA (yellow) which can be later used for quantification of bacterial subpopulations and lozalization. Scale = 20 µm. D) Z position of TasA and MotA expressing cells overtime. Cells expressing TasA (red) are located in the upper region of the colony while MotA expressing cells (yellow) are located in the bottom region. E) 3D representation of the colony growth expressing TasA and MotA. 4 snapshots of the colony growth evolution are shown at different time points (6, 10, 16 and 22 h). TasA cells at 16 and 22 h show the formation of the typical wrinkle found in a *B. subtilis* biofilm. Scale = 20 µm.

#### Concluding remarks

The study of microorganisms at the microscopic level have been pursued since the invention of microscopy in the 17^th^ century. In recent years, the study of bacterial populations and bacterial structures such as biofilms have gained interest and importance. A variety of strategies have been developed to follow these processes over time. At the microscopic level, most existing methods involved a sample processing step before imaging, making it impossible to perform a continuous study over time, and increasing the probability of introducing artefacts into the results.

Recently, some novel strategies have been introduced to improve our ability to image these processes. E. Nadezhdin et al. (29), for example, have developed a new method to study living biofilms using agar segments, thus facilitating bacterial growth and time-lapse microscopy across a cross-section of the biofilm. This approach has allowed the identification of spatial patterns of gene expression focused on the top of the live biofilm, however, the 2D images obtained may be insufficient for the analysis of larger biofilm regions (Table 1).

In this manuscript we have described a new strategy, BacLive, that provides a new method for studying bacterial interactions and living biofilms without sample manipulation. The use of glass-bottomed Petri dishes in combination with fluorescently-labelled bacteria and temperature regulation permits the capture of substantial amounts of data that can be conveniently processed using the BacLive macro, thus reducing the amount of data generated and permitting easier data analysis. This new strategy, in combination with other processing tools, is a value tool for the study of genetic processes in living bacteria, and help toward gaining a more complete perspective into the types of interactions that occur in the environment, as well as aid in applied sciences, for example in the search and development for antimicrobials.

## Materials and Methods

### Strains, media and culture conditions

Routinely, bacterial cells were grown in liquid Lysogeny Broth (LB: 1% tryptone (Oxoid), 0.5% yeast extract (Oxoid) and 0.5% NaCl) medium at 28 °C (*P. carotovorum, B. amyloliquefaciens* FZB42 and *B. subtilis*) with shaking on an orbital platform. Pairwise interaction experiments were performed on LB media while biofilm assays were done on MSgg medium: 100 mM morpholinepropane sulfonic acid (MOPS) (pH 7), 0.5% glycerol, 0.5% glutamate, 5 mM potassium phosphate (pH 7), 50 μg/ml tryptophan, 50 μg/ml phenylalanine, 50 μg/ml threonine, 2 mM MgCl_2_, 700 μM CaCl_2_, 50 μM FeCl_3_, 50 μM MnCl_2_, 2 μM thiamine, 1 μM ZnCl_2_. *Bacillus subtilis* 168 is a domesticated strain used to transform the different constructs into *Bacillus subtilis* NCIB3610. When necessary, antibiotics were added to the media at appropriate concentrations. Strains and plasmids were constructed using standard methods (30).

### Construction of fluorescence labeled strains

For the construction of the double-fluorescently labelled *B. subtilis* strain, P_*tasA*_ was amplified using primers containing EcoRI and XbaI restriction sites and cloned into plasmid ECE756 previously digested with the same restriction enzymes. The resulting phusion was amplified and cloned into pDR183 plasmid. P_*motA*_ was inserted into a pKM003 plasmid. The resulting plasmids were transformed by natural competence into *B. subtilis* 168 replacing the *lacA* and *amyE* neutral locus, respectively. All of the *B. subtilis* strains generated were constructed by transforming *B. subtilis* 168 via its natural competence and then using the positive clones as donors for transferring the constructs into *B. subtilis* NCIB3610 via generalized SPP1 phage transduction (31). Fluorescence labeling plasmid pKM008V was constructed for *B. amyloliquefaciens* FZB42. For that, the P_*veg*_ promoter fragment (300 bp) was extracted from pBS1C3 by digestion with EcoRI and HindIII restriction enzymes, purified and cloned in this case into a pKM003 plasmid, which was previously digested with the same restriction enzymes. pKM008V was then linearized and transformed into *B. amyloliquefaciens* FZB42 by natural competence, and transformants selected by plating on LB agar plates supplemented with spectinomycin (100 μg/ml). *P. carotovorum* was fluorescently labelled by electroporation using the pRL662-gfp kindly donated by Ehr-Min Lai laboratory.

### Time-lapse microscopy

Bacterial interactions and biofilm dynamics in solid medium were visualized by Confocal Laser Scanning Microscopy (CLSM). For biofilm studies, 0.7 µl of the double-labelled *B. subtilis* strain were spotted on MSgg solid media while, for bacterial interaction time-lapse experiments, *B. amyloliquefaciens* FZB42 and *P. carotovorum* labeled strains were spotted at a 0.5 cm distance onto 1.3 mm thick LB agar plates using 35 mm glass bottomed dishes suitable for confocal microscopy (Ibidi). Plates were incubated at 28 °C for 6 h prior to acquisition. Temperature was maintained at 28 °C during the time-course using the integrated microscope incubator. Acquisitions were performed using an inverted Leica SP5 confocal microscope with a 25x NA 0.95 NA IR APO long working distance water immersion objective. Bacterial fluorescence could be visualized from underneath the bottom of the plate and through the agar medium thanks to the long 2.2 mm free working distance of this objective. A special oil immersion medium, Immersol W 2010 (Zeiss) was used instead of water to avoid problems with evaporation during the experiment. Colony fluorescence was followed in multiple regions selected at the start of the experiment, with the acquisition of a series of different focal (z) positions at each region performed automatically at every time-point. Evaporation from the LB agar and its utilization by the growing colonies results in a gradual lowering of the agar surface relative to the objective lens of approximately 250 µm every 24 hours. In order to be able to follow colony dynamics, images were acquired over wide focal range to compensate for the predicted change in colony position during the experiment. Image processing and 3D visualization was performed using ImageJ/FIJI^79, 80^ and Imaris version 7.6 (Bitplane). Expansion rates and distance between colonies were calculated with FIJI, using the change in position of the leading edge of the colony between time-points. Expansion rate results were calculated as an average speed at three different regions of the same colony, with variations expressed as standard deviation while distance was calculated between the closer points of both leading edges of the colonies. To reduce the impact of random vibrations and variations, plotting speed values were smoothed as a floating 4-value average advancing 30 minutes (or 1 time-point) at a time.

## Acknowledgments

We thank Saray Morales Rojas for technical support. This work was supported by grants from ERC Starting Grant (BacBio 637971) and Plan Nacional de I+D+i of Ministerio de Economía y Competitividad and Ministerio de Ciencia e Innovación (AGL2016-78662-R and PID2019-107724GB-I00). C.M.S is funded by the program Juan de la Cierva Incorporación (IJC2018-036923-I). M.V.B.C. and A.I.P.L are funded by the program FPU (FPU17/03874 and FPU19/00289, respectively).

## Supplementary Material

Supplementary Figure 1. Solid media volume and thickness is critical for the correct performance of the experiment. Glass bottomed Petri dishes with different culture media volumes show differences in bacterial growth. 1.3 ml of solid culture media is the best condition tested for the correct bacterial growth without affecting image acquisition.

Supplementary Figure 2. Comparison of strategies for z-slices calculation and acquisition.

A) Brute force strategy is based on capturing all the 3D volume where colonies are expected along the experiment. B) Phased strategy is based on the partial acquisition of 3D volumes according to the day of experiment.

Supplementary Figure 3. BacLive workflow showing the steps performed for the processing of data acquired. Steps with a bold frame are key steps in the BacLive process. Red frames indicate steps repeated in the processing loop.

Supplementary Figure 4. A) Pairwise interaction representation indicating the exact spots used to analyze short-term and long-term stages of the interaction. B) Expansion rates of the FZB42 and *P. carotovorum* leading edges in the short-term stage of the interaction (9 h). The blue line represents the FZB42 leading edge, and the green line represents *P. carotovorum* leading edge.

Supplementary Movie 1. Long term 2D time lapse (18 – 36 h) of the interaction between *P. carotovorum* (green) and FZB42 (cyan).

Supplementary Movie 2. Long term 3D time lapse (18 – 36 h) of the interaction between *P. carotovorum* (green) and FZB42 (cyan). Bacterial colonies are represented as volumes.

Supplementary Movie 3. Long term 3D time lapse (18 – 36 h) of the interaction between *P. carotovorum* (green) and FZB42 (cyan).

Supplementary Movie 4. 3D time lapse of a *B. subtilis* colony leading edge. Spots indicate fluorescent bacteria expressing TasA (red) and MotA (yellow).

Supplementary Movie 5. 3D time lapse of a region of a *B. subtilis* colony where wrinkles are formed. Subpopulations expressing TasA (red) and MotA (yellow) are shown as volumes.

Supplementary Movie 6. 3D time lapse of a region of a *B. subtilis* colony where wrinkles are formed. Subpopulations expressing TasA (red) and MotA (yellow) are shown as volumes. Grey area indicates a region with cells not expressing TasA or MotA.

